# Dietary n-3 PUFA enhances DMI in transition cows by regulating taste transduction gene expression in liver associated with rumen microbial alteration

**DOI:** 10.1101/2023.07.24.550439

**Authors:** Xiaoge Sun, Cheng Guo, Qianqian Wang, Yan Zhang, Zhonghan Wang, Zhijun Cao, Wei Wang, Shengli Li

**Affiliations:** State Key Laboratory of Animal Nutrition, Beijing Engineering Technology Research Center of Raw Milk Quality and Safety Control, College of Animal Science and Technology, China Agricultural University, Beijing, 100193, P. R. China; School of Agriculture, Ningxia University, Yinchuan 750021, China

**Keywords:** PUFA, dry matter intake, bile acid, liver transcriptome, rumen bacterial profile

## Abstract

We hypothesised that the addition of n-3 polyunsaturated fatty acids (PUFAs) in the diet could affect gene expression in the liver and have beneficial effects on the recovery of cows in the transition phase. A total of 30 multiparous non-lactating Holstein dairy cows (35 days before expected calving) were randomly fed a diet with either 1% dry matter (DM) of hydrogenated fatty acid (C16:00 enriched; CON) or 3.5% DM of extruding flaxseed (n-3 enriched; HN3). Parity, body weight (BW), body condition score (BCS) and milk yield were 2.6 ± 1.2, 757.5 ± 65.8 kg, 3.3 ± 0.2 and 10,286.5 ± 1464.8 kg/d (mean ± SD), respectively, at the beginning of the experiment. The relative abundance of *Bacteroidota* (*P* = 0.047) and *Spirochaetota* (*P* = 0.091) was higher and that of *Patescibacteria* (*P* = 0.076) was lower in the HN3 group than in the CON group on prepartum day 4. The DMI of cows was positively correlated with the abundance of bacteria in the rumen (*Spirochaetota*: r = 0.871, *P* < 0.001; *Bacteroidota*: r = 0.896, *P* < 0.001) and the differential expression of genes involved in taste transduction (ACSL1: r = 0.673, *P* < 0.001; PLIN4: r = 0.632, *P* < 0.01; CPT1A: r = 0.694, *P* < 0.001). These results suggest that dietary n-3 PUFA at an appropriate concentration can promote DMI recovery by upregulating the expression of these genes and maintaining the balance of the rumen microbiota.

## Introduction

Dairy cows undergo a physiologically challenging and stressful transition from a pregnant non-lactating stage to a non-pregnant lactating stage owing to parturition and the onset of lactation (Useni et al., 2018). One of the major issues is that cows in the transition phase experience appetite suppression and require 4–6 weeks to increase dry matter intake (DMI) to normal during the post-calving period (Sordillo and Raphael, 2013). Consequently, energy intake is less than that required for milking demands, which is defined as negative energy balance (NEB). During NEB, much more energy should be utilised from tissue storage to support the energy requirement, and the increased lipid mobilisation compromises liver function (Herdt, 2000).

Lipid mobilisation alters the concentration and composition of plasma non-esterified fatty acids (NEFAs), which are metabolic biomarkers of NEB. The effects of NEFAs are related to the occurrence of clinical ketosis, abomasum displacement and endometritis, all of which can increase the risk of animal removal (LeBlanc, 2010, Puppel and Kuczyńska, 2016). Saturated (e.g. palmitic and stearic acids) and monosaturated (e.g. oleic acid) fatty acids are the predominant fats found in plasma at the transition phase; however, the concentration of polyunsaturated fatty acids (PUFA) such as n-3 fatty acids is low during this period (Sordillo and Raphael, 2013).

n-3 PUFAs have several beneficial effects on human health, especially in metabolic diseases (Simopoulos, 2002). Therefore, strategies for enhancing n-3 PUFA concentration in milk have attracted substantial attention. An easily and widely available source of n-3 PUFAs (α-linolenic acid, C18:3n-3) is flaxseed. Previous studies have comprehensively elucidated the effects of n-3 PUFA supplementation on milk production and milk FA profile (Gonthier et al., 2005, Akraim et al., 2007, Petit et al., 2007). Flaxseed is effective in increasing n-3 PUFA content in plasma and milk fat. However, the effects of dietary n-3 PUFAs on DMI, bile acid, rumen microbiota or hepatic gene expression profile during the transition period of dairy cows remain unclear. Petit et al. (2007) reported that compared with supplementation of saturated fats, supplementation of whole flaxseed (3.3 and 11.0% of DM prepartum and postpartum, respectively) beginning 6 weeks before calving increased the concentration of liver glycogen and decreased the concentration of triglycerides in multiparous cows. Therefore, feeding flaxseed to pre-calving cows may be a useful strategy for preventing the development of fatty liver in dairy cows undergoing the transition phase (Petit et al., 2007). We hypothesise that dietary supplementation with n-3 PUFAs affects the hepatic gene expression profile and has beneficial effects on the recovery of cows after the transition period.

This study aimed to evaluate the effects of dietary n-3 PUFAs on rumen fermentation and microbiome, bile acids in the liver and hepatic gene expression profile associated with alterations in DMI during the pre- and post-calving periods.

## Materials and methods

Animals involved in this study were maintained according to the guidelines recommended by the China Agricultural University Laboratory Animal Welfare and Animal Experimental Ethical Inspection Committee. All procedures involving animals were reviewed and approved by the Animal Experimental Ethical Inspection Committee.

## Animals and experimental design

All animal experiments were conducted at Beijing Zhongdi Animal Husbandry Technology Co., LTD. A total of 30 multiparous non-lactating Holstein dairy cows (35 d before expected calving) were housed in a free-stall barn (with a rubber bed and rice hull bedding) equipped with the Roughage Intake Control (RIC) system (INSENTEC, Marknesse, Netherlands) for a 7-d adaptation period followed by a 28-d prepartum and a 28-d postpartum period. During the experiment, cows were fed either an isoenergetic and isoprotein diet in a randomised manner: CON group, 1% dry matter (DM) of hydrogenated fatty acid (C16:00 enriched) or HN3 group, 3.5% DM of extruding flaxseed (n-3 enriched). Six cows were removed at various stages of the experiment for the following reasons: early calving (1/6 cows), pneumonia (1/6 cows), displacement of the abomasum (2/6 cows) and lameness (2/6 cows). Eventually, the experimental data of 24 cows (CON, n = 12; HN3, n = 12) were used for statistical analysis. Parity, body weight (BW), body condition score (BCS) and milk yield were 2.6 ± 1.2, 757.5 ± 65.8 kg, 3.3 ± 0.2 and 10,286.5 ± 1464.8 kg/previous lactation (mean ± SD), respectively, at the beginning of the experiment. Cows were offered a close-up total mix ration (TMR) before calving, followed by a postpartum milk cow TMR (Table 1). Diet was delivered to cows twice a day at 0700 and 1500 h and cows had free access to TMR.

**Table 1.** Ingredients and chemical composition of TMR (DM basis) for close-up and lactating cows.

| Item | Prepartum |  | Postpartum |  |
| --- | --- | --- | --- | --- |
|  | CON | HN3 | CON | HN3 |
| Ingredients (%) |  |  |  |  |
| Oat hay | 47.82 | 47.62 |  |  |
| Alfalfa hay |  |  | 10.35 | 10.24 |
| Corn silage | 21.70 | 21.61 | 30.43 | 30.10 |
| Alfalfa silage |  |  | 7.84 | 7.76 |
| Corn fine | 3.88 | 2.74 | 9.46 | 9.37 |
| Soybean Hulls | 7.72 | 7.70 | 2.65 | 2.63 |
| Corn steam flakes |  |  | 10.32 | 10.21 |
| Soybean meal | 7.61 | 6.58 | 16.15 | 14.48 |
| Corn gluten feed | 4.08 | 4.06 |  |  |
| Urea | 0.61 | 0.61 |  |  |
| Cottonseed fuzzy |  |  | 5.36 | 5.33 |
| Sugarcane molasses |  |  | 2.59 | 2.56 |
| Diamond V XP | 0.26 | 0.26 | 0.16 | 0.16 |
| Hydrogenated FA | 1.00 |  | 1.00 |  |
| Extruded soybean |  |  |  |  |
| Extrude flaxseed |  | 3.50 |  | 3.50 |
| Dry cow premix | 4.31 | 4.31 |  |  |
| DCAD supplement | 1.01 | 1.01 |  |  |
| Milking cow premix |  |  | 3.69 | 3.66 |
| Nutrition level |  |  |  |  |
| DM(kg) | 12.97 | 13.02 | 17.27 | 17.26 |
| CP(%) | 15.56 | 15.52 | 17.38 | 17.39 |
| NE<sub>L</sub>(Mcal/kg) | 1.27 | 1.28 | 1.68 | 1.71 |
| NFC(%) | 30.36 | 29.30 | 41.77 | 41.35 |
| NDF(%) | 49.07 | 49.67 | 30.39 | 31.64 |
| EE(%) | 3.78 | 3.82 | 5.43 | 5.49 |
| C16:0 (g/d) | 91.13 | 43.36 | 196.21 | 94.02 |
| n-6(g/d) | 83.53 | 87.91 | 191.60 | 226.57 |
| n-3(g/d) | 49.63 | 136.52 | 39.33 | 142.56 |
| n-6: n-3 | 1.68: 1 | 0.64: 1 | 4.87: 1 | 1.59: 1 |

DM: Dry matter, CP: Crude protein, NE<sub>L</sub>: Net energy of lactation, a calculated value according to NRC (2001), NFC: Non-fibrous carbohydrate, NDF: Neutral detergent fiber, EE: Ether extract, UFA: Unsaturated fat acid, n-6: C18:2n6 (Linoleic acid, LA), n-3: C18:3n3 ( $\alpha$ - Linolenic acid, ALA). Dry cow premix: 1 kg of premix included vitamin A 918,750 IU, vitamin D3 253,125 IU, vitamin E 8,066 mg, niacin 6,650 mg, Ca 35 g, P 20 g, Cu 1146mg, Mn 4248 mg, Zn 5362 mg, I 45 mg, Se 36 mg. Milking cow premix: 1 kg of premix included vitamin A 440,000 IU, vitamin D3 110,000 IU, vitamin E 4000 IU, niacin 400 mg, Ca 152 g, P 41 g, Cu 750 mg, Mn 1140 mg, Zn 2970 mg, I 30 mg, Se 24 mg. DCAD supplement: - $107.43 \text{ (mEq/kg)} = (\text{Na}\%/0.023 + \text{K}\%/0.039 - \text{Cl}\%/0.0355 - \text{S}\%/0.016)$ . CON: Control treatment, HN6: High n-6 polyunsaturated fatty acid (PUFA) treatment, HN3: High n-3 PUFA treatment.

## Data and sample collection

Feed intake was monitored daily using the RIC system (INSENTEC, Marknesse, the Netherlands). Samples of TMR and orts were collected at weeks –4, –3, –2 and –1 before calving and weeks 1, 2, 3 and 4 weeks after calving and stored at -20°C for chemical composition analysis at the State of Key Laboratory of Animal Nutrition in China. Milk production was recorded at each milking session using the ALPROTM system (DeLaval©, Tumba, Sweden).

Blood samples were collected from the tail vein via venepuncture of the coccygeal vessels 2 h after morning feeding at 0900 h before 4 d of calving and after 28 d of calving. Samples were collected in 10-mL vacutainer tubes (Vacutainer, Becton Dickinson, Rutherford, NJ). The tubes were centrifuged at 3,000 ×g for 12 min at 4°C. Serum was collected and transferred to 2-mL plastic scintillation vials and stored at –20°C until further analysis.

Before 4 d of calving and after 28 d of calving, 100-mL rumen fluid was collected from each cow through a stomach tube as described previously (Shen et al., 2012). The rumen fluid was immediately filtered using four layers of gauze, and the filtrate was divided into two 50-mL centrifuge tubes. One of the aliquots was used to measure pH using sophisticated handheld pH meters (Starter 300; Ohaus Instruments Co., Ltd., Shanghai, China), whereas the other aliquot was stored at –20℃ and used for analysing volatile fatty acids (VFAs), ammonia (NH3-N) and microbial proteins (MCPs).

Liver tissue samples were collected from 6 cows in each group at two stages: 4 d before calving and 28 d after calving. The samples were collected via blind percutaneous needle biopsy under local anaesthesia, immediately placed in liquid nitrogen and stored at −80°C.

## Liver biopsy

Liver biopsy was performed between 0900 and 1100 h before 4 d and after 28 d of calving according to a method described by Casey et al. (2021). An area of 15–20 cm^2^ over the 11th and 12th intercostal spaces on a line from the hip joint to the elbow was clipped. For sterilisation, the site was washed with soapy water, shaved, scrubbed thrice with a betadine solution and swabbed with 70% ethanol. Subsequently, 50 mL of lidocaine was subcutaneously injected at the 11th and 12th intercostal spaces, and a 1-cm incision was created using a scalpel blade after the site was numb to touch. A sterile, 9-mm custom biopsy needle (similar to a trocar cannula) was inserted through the peritoneal tissue to the liver. The stylus was retracted, and the instrument was inserted 2–3 cm into the liver, which caused a slight vacuum within the shaft of the instrument to recover the biopsy sample. Approximately 3 g (wet weight) of liver tissue was collected and divided into three tubes. The tissue samples were immediately snap-frozen in liquid nitrogen and stored at –80℃ for further analysis. The incision was closed using an absorbable suture (#2 Supramid; Jinhuan Medical Products Co., LTD, Shanghai, China). Incision sites were monitored and disinfected with betadine solution at least once daily for 7 days.

## Chemical analysis

The DM content of feeds/TMR was determined at 135°C for 2 h (AOAC, 1990). The crude protein (CP) content of feed/TMR was determined (AOAC, 1990) using an Elementar Rapid N Exceed system (Elementar, Germany). The concentration of ether extracts (EEs) was measured according to method 920.39 of the Association of Official Analytical Chemists (AOAC, 1990). Feed samples were analysed for acid detergent fibre (ADF) (AOAC, 1990) and neutral detergent fibre (NDF) (Van Soest et al., 1991) using the ANKOM 2000i automatic fibre analyser (Beijing Anke Borui Technology Co., Ltd., Beijing, China). The methylation of dietary fats was assessed using a method described by Loor and Herbein (2001). The concentration of FA methyl esters was determined on a gas chromatography system (Agilent GC6890, US) equipped with a flame ionisation detector and a fused silica capillary column (DB-23, 60.0 m×0.25 mm×0.25 μm, USA).

The concentration of orexins, peptide tyrosine–tyrosine (PYY) and neuropeptide Y (NPY) was evaluated on an enzyme-labelled instrument (Thermo Multiskan Ascent, America) based on enzyme-linked immunosorbent assay (ELISA).

The composition and concentration of bile acids (BAs) were analysed through ultra-performance liquid chromatography–tandem mass spectrometry (LC-MS/MS) with modifications in a method described previously (Yin et al., 2017). The full names and respective abbreviations of different BAs are shown in Table S1. BAs were extracted from serum as follows: 50 μL of serum was mixed with 10 μL of internal standards (Sigma-Aldrich, St. Louis, USA). Thereafter, 240 μL of cold methanol was added, and the mixed solution was vortexed thrice for 10 s each time and incubated at –20°C for 20 min. The solution was centrifuged at 10,000 g for 10 min at 4°C. The supernatant was transferred to an autosampler vial and was dried with nitrogen. The dry residue was reconstituted by adding 100 μL of methanol/water (1:1). Finally, the redissolved solution was centrifuged at 10,000 g for 5 min at 4°C, and the supernatant was transferred into another autosampler vial for LC-MS/MS analysis. BAs were extracted from the liver as follows: 20 mg of liver tissue was mixed with 10 μL of internal standards (Sigma-Aldrich, St. Louis, USA). Subsequently, 450 μL of cold methanol was added, and the mixed solution was vortexed for 60 s. The solution was centrifuged at 10,000 g for 10 min at 4°C, and the supernatant was transferred to an autosampler vial. Thereafter, 450 μL of cold methanol was added for the second extraction, and the sample was centrifuged. The supernatants from the two extractions were combined and diluted 10 times with methanol–water (1:1) for LC-MS/MS analysis.

An Agilent 1290 ultra-performance liquid chromatography system coupled with a 6490 triple quadrupole mass spectrometer (Agilent Techonologies Inc, CA, USA) was used for the quantitative analysis of BAs. The Acquity UPLC BEH C18 column (1.7 μm, 2.1×100 mm, Waters) was used for separating BAs at 65°C. Mass spectrometry was performed with the negative ion mode. Various isotope-labelled BAs (Sigma-Aldrich, St. Louis, USA) were used as an internal standard to resolve errors arising owing to sample pre-treatment and mass spectrometry analysis. The content of BAs in the liver and serum was originally expressed as ng/mL and ng/g, respectively. Subsequently, the data were visualised and analysed in the log base 2 (log2) form.

The concentration of ammonia and MCPs was determined via spectrophotometry as described in previous studies (Verdouw et al., 1978, Cui et al., 2016). The concentration of VFAs was measured via gas chromatography (6890N; Agilent technologies, Avondale, PA, USA) as described previously (Zhang and Yang, 2011).

## DNA extraction and determination

Total DNA was extracted from rumen fluid using the TIANGEN® TIANamp Stool DNA Kit (Tiangen Biotech Co., Ltd., Beijing, China). The concentration and purity of the extracted DNA were determined using the NanoDrop-1000 Spectrophotometer (Thermo Fisher Scientific, Waltham, MA, USA). The V3–V4 region of the 16S rRNA gene was amplified according to a method described previously (Hao et al., 2020). The forward primer sequence 341F (CCTAC-GGGNGGCWGCAG) and reverse primer sequence 806R (GGACTACHVGGGTATCTAAT) were used for amplification (Guo et al., 2017). Amplicons were extracted from 2% agarose gels, purified using the AxyPrepDNA Gel Extraction Kit (Axygen Biosciences, Union City, CA, USA) and quantified on the GeneAmp PCR System 9700 (Applied Bio-systems, Foster City, CA, USA).

Purified amplicons were pooled in equimolar amounts and sequenced using the Miseq PE300 platform (Illumina, Inc., San Diego, CA, USA) to generate paired-end (2 × 300) sequences. The original data were stored in the Fastq format and processed for quality control using the Trimmomatic (version 0.36) (Bolger et al., 2014) and Pear (version 0.9.6) (Zhang et al., 2014) software. In the Trimmomatic software, a sliding window strategy was adopted, with the window size set to 50 bp, the average base quality score set to 20 and the minimum reserved sequence length set to 120. The Pear software was used to remove Ń sequences. Reads with unknown nucleotides (>10%) were removed, and paired-end clean tags were merged as raw tags using the FLASH (version 1.2.11) software as described by Magoč and Salzberg (2011). The minimum overlap was set to 10 bp and the mismatch rate was set to 0.1 to obtain Fasta sequences. Chimeric Fasta sequences were removed using UCHIME (Shokralla et al., 2015), and a *de novo* method was used to remove undesirably short sequences. A total of 2,467,080 raw tags and 2,418,713 clean tags were obtained. An average of 94,887 raw tags and 93,027 clean tags were obtained for each sample. After removing noise, high-quality tags were clustered into operational taxonomic units (OTUs) with a clustering threshold of ≥97% identity using the Vsearch (version 2.7.1) software according to the UPARSE pipeline (Edgar, 2013). Finally, 735,982 tags and 52,178 OTUs were obtained. An average of 28,307 tags and 2,006 OTUs were obtained for each sample. Representative sequences (those with the highest relative abundance in each cluster) were classified using BLAST (version 2.6.0) (Camacho et al., 2009), which is based on the SILVA (version 138.1) database (https://www.arb-silva.de/) (Pruesse et al., 2007). The abundance of each taxonomy was visualised using the Krona (version 2.6) software (Ondov et al., 2011). Alpha diversity indices (e.g. Chao1, observed species and Shannon indices) were evaluated using QIIME 2 (Hall and Beiko, 2018), and sequences were aligned using MAFFT (version 7.0) (Katoh and Standley, 2013).

## RNA extraction from liver tissues

Total RNA was extracted from 50–60 mg of frozen bovine liver tissues (n = 12) using the guanidinium thiocyanate method (Pareek et al., 2019) (TRIzol reagent; Thermo Fisher Scientific Inc., Waltham, MA, USA). The extracted RNA was purified to remove genomic DNA contamination using the RNase-free DNase clean-up kit (Thermo Fisher Scientific Inc., Waltham, MA, USA). The degradation and contamination of RNA were monitored on 1% agarose gels, and its concentration and purity were measured on a NanoDrop spectrophotometer (Thermo Scientific, DE, USA). RNA integrity was analysed on the Agilent 2100 Bioanalyzer (Agilent Technologies, CA, USA).

## Library preparation for transcriptomic sequencing

The RNA integrity number of all biological samples (n = 12) ranged from 6.9 to 8.5. A total amount of 1.5-μg RNA per sample was used as an input for RNA sample preparation, and two biological replicates were prepared for each biological sample. Sequencing libraries were generated using the NEBNext® Ultra™ RNA Library Prep Kit for Illumina® (NEB, USA) following the manufacturer’s instructions, and index codes were added to attribute sequences to each sample. The library fragments were purified on the AMPure XP system (Beckman Coulter, Beverly, USA) to select cDNA fragments of 200–250 bp in length. The size-selected, adaptor-ligated cDNA sample was incubated with 3 μL of USER Enzyme (NEB, USA) for 15 min at 37°C, followed by 5 min at 95°C. Subsequently, PCR was performed using Phusion High-Fidelity DNA polymerase, Universal PCR primers and Index (X) Primer. PCR products were purified (AMPure XP system), and their quality was assessed on the Agilent 2100 Bioanalyzer system. The prepared library was sequenced on the Illumina Novaseq 6000 platform by the Beijing Allwegene Technology Co., LTD. (Beijing, China), and 150-bp paired-end reads were generated.

## Quantitative real-time polymerase chain reaction (qPCR)

The differentially expressed genes (DEGs) involved in the taste transduction (ACSL1, CPTIA, PLIN4) upregulated in HN3 treatment were selected to do validation using quantitative real-time polymerase chain reaction (qPCR). Glyceraldehyde-3-phosphate dehydrogenase (GAPDH) as a housekeeping gene. Primers were designed to amplify specific cDNA sequences in each target gene using NCBI website tool (https://www.ncbi.nlm.nih.gov/tools/primer-blast). Sequences are summarized in Table S2. To avoid possible amplification of contaminating genomic DNA, all RNA samples were treated with DNase before cDNA synthesis.

Reverse transcription reactions were conducted using 1µg of total RNA as a template, 4µL 5×FastKing-RT SuperMix (Tiangen Biotech, Beijing, China), add RNase-Free ddH_2_O to a final volume of 20 μL.

Real-time PCR reactions were performed in triplicate in 96-well plates using SYBR Green I universal master mix (Applied Biosystems), 500 nM each of the forward and reverse primers (Integrated DNA Technologies, Coralville, IA), and 1 μL of diluted cDNA in a final reaction volume of 20 μL per well. Amplification and data collection were carried out using an applied Biosystems QuantStudio 6&7 instrument with reaction conditions of 15 min at 95°C, and 40 cycles of 10 s at 95°C and 32 s at 60°C.

## Statistical analysis

The daily mean DMI and milk production during the pre- and post-calving periods were evaluated using the two-sample t-test in the SAS (version 9.4) software (SAS Institute Inc., Cary, NC). The 2^-ΔΔCT^ method was used as a relative quantification strategy for qPCR data analysis (Livak and Schmittgen, 2001).

The parameters of rumen fluid fermentation (i.e. pH and the levels of NH3-N, MCPs, BAs, acetate, propionate, butyrate and total VFAs), bacterial alpha diversity indices (e.g. Chao1, observed species and Shannon indices), qPCR data and concentration of serum metabolites (e.g. orexins, PYY, LPS and NPY) were analysed using the MIXED procedure in SAS. Relevant model were constructed using the following equation (1):

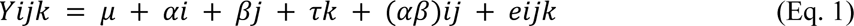

In the abovementioned equation, Yijk represents the dependent variable, μ represents the overall mean, αi represents the fixed effect of treatment (I = 1, 2), βj represents the fixed effect of sampling time (j = –4 d, 28 d), τk represents the random effect of cows, (αβ)ij represents the interaction between treatment and sampling time and eijk represents the residual error. The log2 data of BA concentration were analysed using Eq. 1.

The Kruskal–Wallis test was used to estimate differences in the number of sequenced reads of rumen microbiota between samples. The test was implemented using the *qvalue* package in the R software (version 2.11.1). Differences in the abundance of reads were tested using Metastats (White et al., 2009).

The Wald test was used to generate nominal and adjusted *P*-values and log2 fold changes to identify differentially expressed genes (DEGs) in liver tissue (Love et al., 2014). Genes with *P*-values of <0.05 and |log2 fold change| values of >2 were considered differentially expressed. Spearman correlation coefficients were evaluated using the *pheatmap* package in R to examine the correlation between different parameters and DEG expression.

A *P*-value of <0.05 indicated significant differences, and a tendency towards significance was considered at 0.05 ≤ *P* < 0.10.

## Results

### Milk production and dry matter intake

The data on milk production and DMI are summarised in Table 2. HN3 supplementation significantly increased milk production during the post-calving period (*P* < 0.05) and increased DMI during both pre-calving (*P* < 0.01) and post-calving (*P* < 0.05) periods.

**Table 2.** Milk production and DMI during the pre- and post-calving periods.

| Item | CON | HN3 | SEM | *P*-Value |
| --- | --- | --- | --- | --- |
| Milk production (kg/d) | 35.54 | 41.79 | 1.01 | <0.01 |
| Prepartum DMI (kg/d) | 11.58 | 12.75 | 0.17 | <0.01 |
| Postpartum DMI (kg/d) | 17.81 | 20.26 | 0.43 | <0.01 |

DMI, dry matter intake; CON, group of cows fed a control diet; HN3, group of cows fed a high-concentration n-3 PUFA-rich diet; SEM, standard error of the mean.

### Feeding factor in serum

The concentration of orexins, LPS and NPY in serum was higher in the HN3 group than in the CON group throughout the experiment (Table 3; *P* < 0.05). During the transition period, the concentration of PYY in serum was higher in the HN3 group than in the CON group (*P* < 0.05). However, the concentration of PYY in serum did not differ between the HN3 and CON groups during the pre- or post-calving periods.

**Table 3.** Effect of n-3 PUFA on the concentration of orexins, PYY, NPY and LPS in the serum of cows during the transition period.

| Item | Treatment |  | Time |  | SEM | *P*-value |  |  |
| --- | --- | --- | --- | --- | --- | --- | --- | --- |
|  | CON | HN3 | Pre4d | Post28d |  | Trt | Time | Trt×Time |
| Orexins(ng/mL) | 1.60 | 1.88 | 1.74 | 1.71 | 0.069 | <0.01 | 0.57 | 0.56 |
| PYY(pmol/L) | 7.45 | 8.17 | 8.15 | 7.48 | 0.307 | 0.15 | 0.50 | 0.42 |
| NPY(pg/mL) | 111 | 132 | 120 | 122 | 5.95 | <0.01 | 0.72 | 0.08 |
| LPS(U/L) | 37.4 | 42.9 | 42.1 | 38.2 | 2.45 | 0.03 | 0.05 | 0.52 |

PYY, peptide tyrosine-tyrosine; NPY, neuropeptide Y; LPS, Lipopolysaccharide. Pre4d, 4 days before calving; Pos28d, 28 days after calving; SEM, standard error of the mean.

### Rumen fermentation parameters

The pH was not affected by any dietary supplement during the pre- and post-calving periods (Table 4). The concentration of NH3-N was higher in the HN3 group than in the CON group (*P* < 0.05). Moreover, the concentration of NH3-N was significantly lower on day 28 after calving than on day 4 before calving (*P* < 0.05). The concentration of MCPs was not different between the two groups during the pre- and post-calving periods. There was a tendency increase for the concentration of acetate, butyrate and total VFA in the HN3 group than in the CON group (*P* < 0.1). The concentration of acetate and the ratio of acetate-to-propionate decreased from day 4 before calving to day 28 after calving (*P* < 0.05). In addition, the ratio of acetate-to-propionate was not different between the two groups during the pre- and post-calving periods but decreased from day 4 before calving to day 28 after calving (*P* < 0.01).

**Table 4.**
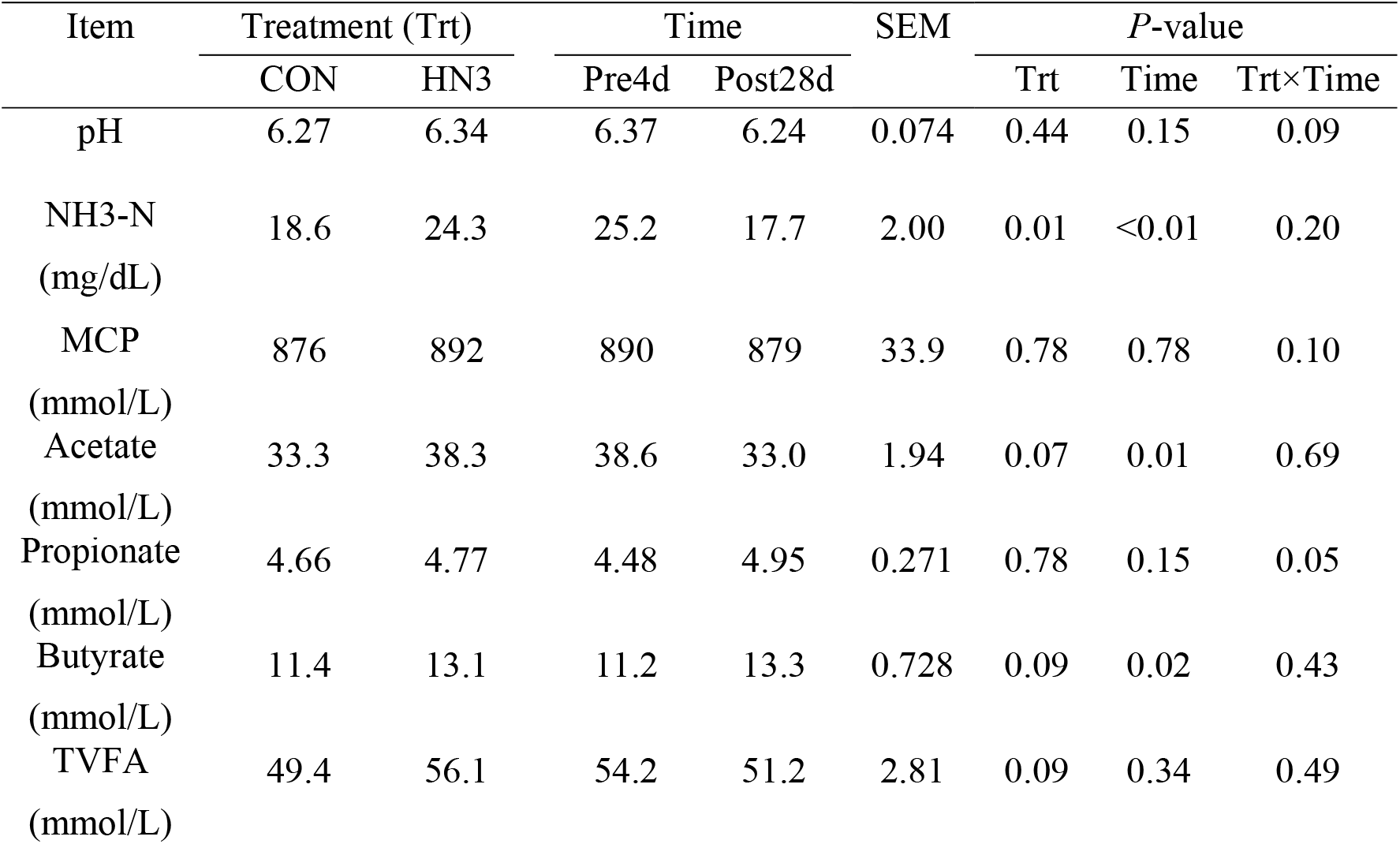
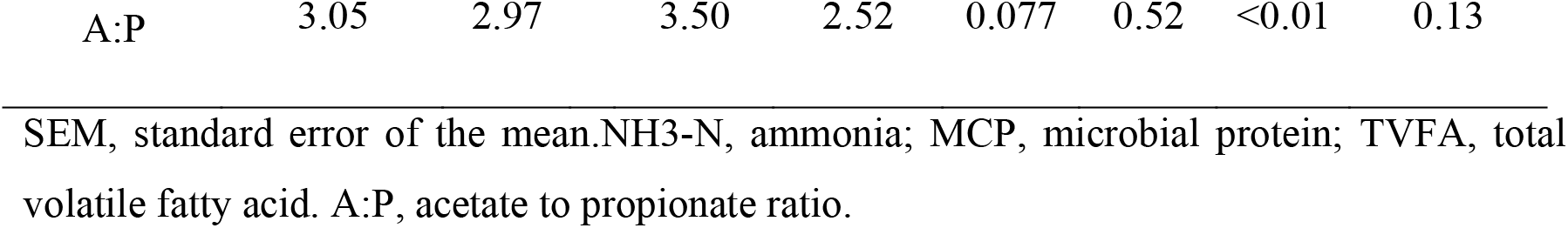
Effects of n-3 PUFA on fermentation parameters in the rumen fluid of cows during the pre- and post-calving periods.

| Item | Treatment (Trt) |  | Time |  | SEM | P-value |  |  |
| --- | --- | --- | --- | --- | --- | --- | --- | --- |
|  | CON | HN3 | Pre4d | Post28d |  | Trt | Time | Trt×Time |
| pH | 6.27 | 6.34 | 6.37 | 6.24 | 0.074 | 0.44 | 0.15 | 0.09 |
| NH<sub>3</sub>-N<br>(mg/dL) | 18.6 | 24.3 | 25.2 | 17.7 | 2.00 | 0.01 | <0.01 | 0.20 |
| MCP<br>(mmol/L) | 876 | 892 | 890 | 879 | 33.9 | 0.78 | 0.78 | 0.10 |
| Acetate<br>(mmol/L) | 33.3 | 38.3 | 38.6 | 33.0 | 1.94 | 0.07 | 0.01 | 0.69 |
| Propionate<br>(mmol/L) | 4.66 | 4.77 | 4.48 | 4.95 | 0.271 | 0.78 | 0.15 | 0.05 |
| Butyrate<br>(mmol/L) | 11.4 | 13.1 | 11.2 | 13.3 | 0.728 | 0.09 | 0.02 | 0.43 |
| TVFA<br>(mmol/L) | 49.4 | 56.1 | 54.2 | 51.2 | 2.81 | 0.09 | 0.34 | 0.49 |
| A:P | 3.05 | 2.97 | 3.50 | 2.52 | 0.077 | 0.52 | <0.01 | 0.13 |

SEM, standard error of the mean.NH3-N, ammonia; MCP, microbial protein; TVFA, total volatile fatty acid. A:P, acetate to propionate ratio.

### Rumen microbial profile

#### Effect of diet on the rumen microbial profile

The Chao1, observed species and Shannon indices did not differ between the HN3 and CON groups during the pre- and post-calving periods (Figure 1).

**Figure 1.**
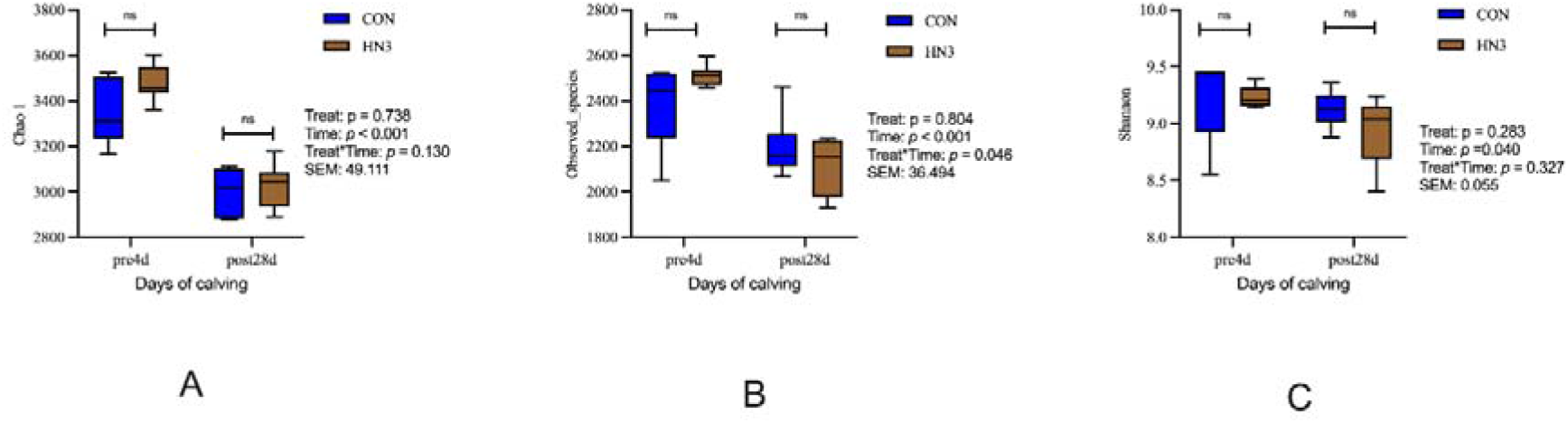
Effects of n-3 PUFA on bacterial diversity and richness in the rumen of cows during the pre- and post-calving periods. CON, group of cows fed a control diet; HN3, group of cows fed a high-concentration n-3 PUFA-rich diet. (A) Chao1 index; (B) Number of observed species; (C) Shannon index. SEM, standard error of the mean; ns, no significant difference between the groups.

At the phylum level, the most abundant bacteria were *Bacteroidota*, *Firmicutes*, *Proteobacteria*, *Actinobacteriota*, *Patescibacteria* and *Spirochaetota* (Figure 1S). The relative abundance of *Bacteroidota* (*P* < 0.05) and *Spirochaetota* (*P* = 0.09) was higher and that of *Patescibacteria* (*P* = 0.07) was lower in the HN3 group than in the CON group on day 4 before calving (Table 5). During the post-calving period, the relative abundance of *Spirochaetota* (*P* < 0.05) and *Elusimicrobia* (*P* = 0.06) were higher in the HN3 group than in the CON group.

**Table 5.** Relative abundance of microorganisms at the phylum level (> 0.1%).

| Phylum (%) | Time | Treatment |  |  | *P*-value (treatment)<br>CON vs. HN3 | *P*-value (time) |  |
| --- | --- | --- | --- | --- | --- | --- | --- |
|  |  | CON | HN3 | SEM |  | Pre4_CON vs.<br>post28d_CON | Pre4_HN3 vs.<br>post28d_HN3 |
| *Spirochaetota* | Pre4d | 0.40 | 0.56 | 0.028 | 0.09 | <0.01 | 0.01 |
|  | Post28d | 1.04 | 1.44 | 0.075 | 0.04 |  |  |
| *Desulfobacterota* | Pre4d | 0.57 | 0.54 | 0.016 | 0.77 | 0.04 | 0.05 |
|  | Post28d | 0.35 | 0.40 | 0.018 | 0.49 |  |  |
| *Firmicutes* | Pre4d | 44.60 | 39.49 | 1.085 | 0.16 | 0.49 | 0.58 |
|  | Post28d | 41.89 | 37.10 | 1.169 | 0.32 |  |  |
| *Elusimicrobia* | Pre4d | 0.08 | 0.17 | 0.015 | 0.10 | 0.19 | 0.19 |
|  | Post28d | 0.04 | 0.08 | 0.008 | 0.06 |  |  |
| *Bacteroidota* | Pre4d | 46.34 | 52.80 | 1.149 | 0.04 | 0.18 | 0.71 |
|  | Post28d | 50.66 | 54.18 | 1.027 | 0.41 |  |  |
| WPS-2 | Pre4d | 0.07 | 0.10 | 0.005 | 0.20 | 0.02 | 0.01 |
|  | Post28d | 0.01 | 0.04 | 0.007 | 0.18 |  |  |
| *Patescibacteria* | Pre4d | 1.65 | 1.25 | 0.042 | 0.07 | 0.01 | 0.48 |
|  | Post28d | 1.06 | 1.12 | 0.052 | 0.74 |  |  |
| *Cyanobacteria* | Pre4d | 0.31 | 0.45 | 0.023 | 0.21 | 0.97 | 0.55 |
|  | Post28d | 0.31 | 0.38 | 0.032 | 0.49 |  |  |
| *Actinobacteriota* | Pre4d | 2.05 | 2.83 | 0.190 | 0.42 | 0.04 | <0.01 |
|  | Post28d | 0.49 | 0.52 | 0.028 | 0.76 |  |  |
| *Fibrobacterota* | Pre4d | 0.44 | 0.38 | 0.021 | 0.62 | 0.26 | 0.26 |
|  | Post28d | 0.33 | 0.53 | 0.038 | 0.10 |  |  |

CON, group of cows fed a control diet; HN3, group of cows fed a high-concentration n-3 PUFA-rich diet.

#### Effect of time on the rumen microbial profile

The Chao1 (*P* < 0.001), observed species (*P* < 0.001) and Shannon (*P* < 0.05) indices were lower on day 28 after calving than on day 4 before calving in the HN3 and CON groups (Figure 1).

The abundance of *Spirochaetota* was 2-fold higher (*P <* 0.01), that of *Candidatus Eremiobacterota* (WPS-2) was 2-fold lower (*P* < 0.05) and that of *Actinobacteriota* was 3-fold lower (*P* < 0.05) on day 28 after calving than on day 4 before calving (Table 5).

### Bile acids in the liver and serum

#### Effect of diet on the concentration of bile acids

The concentration of most BAs in serum was not affected by the diet (Table 6). However, that of TCDCA (*P* = 0.059) and TDCA (*P* = 0.061) was higher in the HN3 group than in the CON group on day 4 before calving (Figure S2). Similarly, the concentration of most BAs in the liver was not affected by the diet (Table 7), except LCA concentration was lower and HDCA concentration was higher (*P* < 0.05) in the HN3 group than in the CON group on day 28 after calving (Figure 3S).

**Figure 2.**
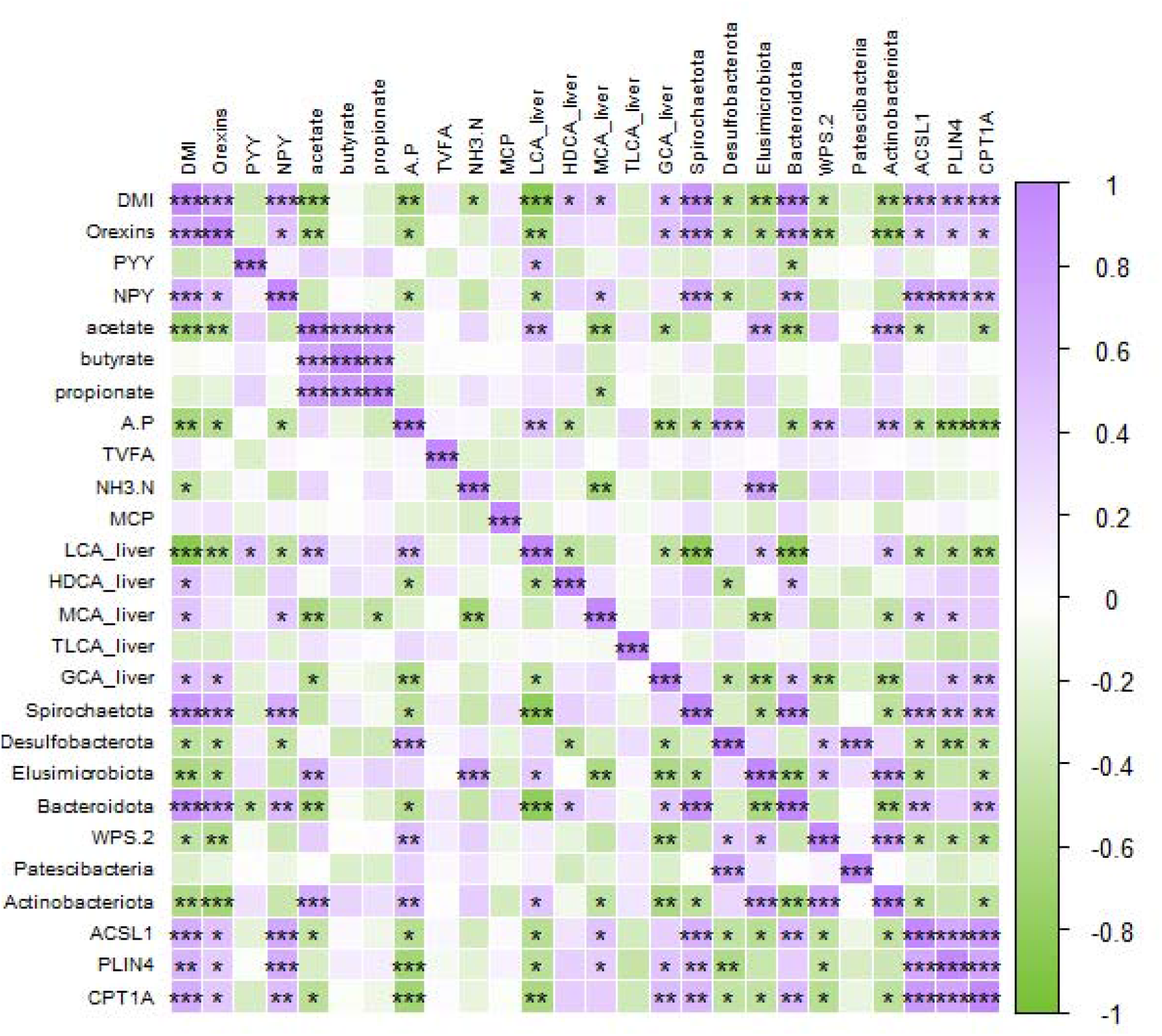
Correlation of DMI with the concentration of different metabolites, abundance of bacteria in the rumen and expression of differentially expressed genes. DMI, dry matter intake; PYY, peptide tyrosine–tyrosine; NPY, neuropeptide Y; A:P, acetate-to-propionate ratio; NH_3_-N, ammonia; TVFA, total volatile fatty acid; MCP, microbial crude protein; LCA_liver, lithocholic acid in the liver; HDCA, hyodeoxycholic acid; MCA_liver, muricholic acid in the liver; TLCA_liver, taurolithocholic acid in the liver; GCA_liver, glycocholic acid in the liver. ACSL1, PLIN4 and CPT1A were identified as differentially expressed genes (present in liver tissue) between the CON and HN3 groups (*, P < 0.05; **, P < 0.01; ***, P < 0.001).

**Figure 3.**
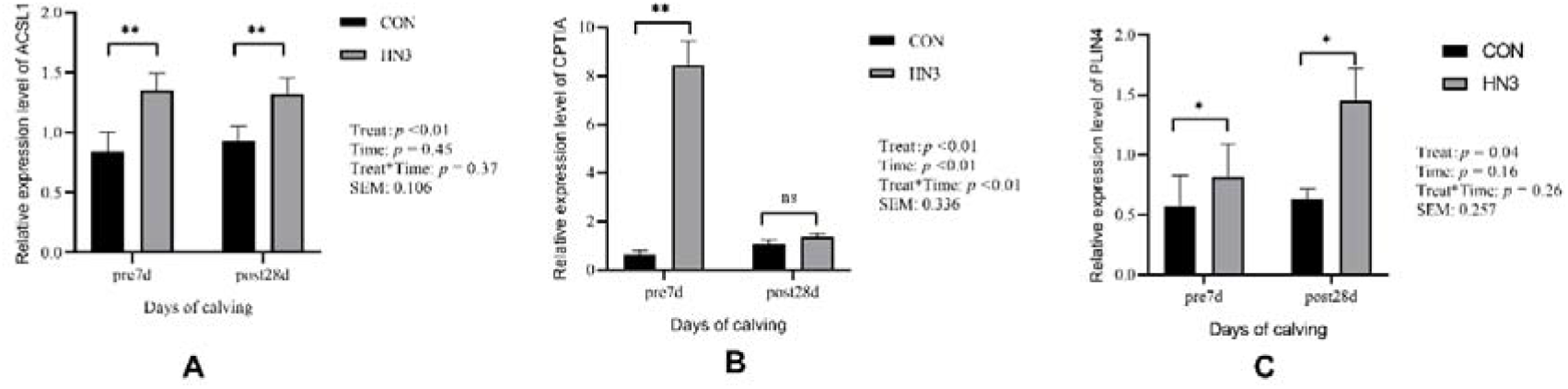
Effects of n-3 PUFA on gene expression using QPCR validation. CON, group of cows fed a control diet; HN3, group of cows fed a high-concentration n-3 PUFA-rich diet. (A) ACSL1 gene expression; (B) ACPT1A gene expression; (C) PLIN4 gene expression. SEM, standard error of the mean; *, P < 0.05; **, P < 0.01; ns, no significant difference between the groups.

**Table 6.**
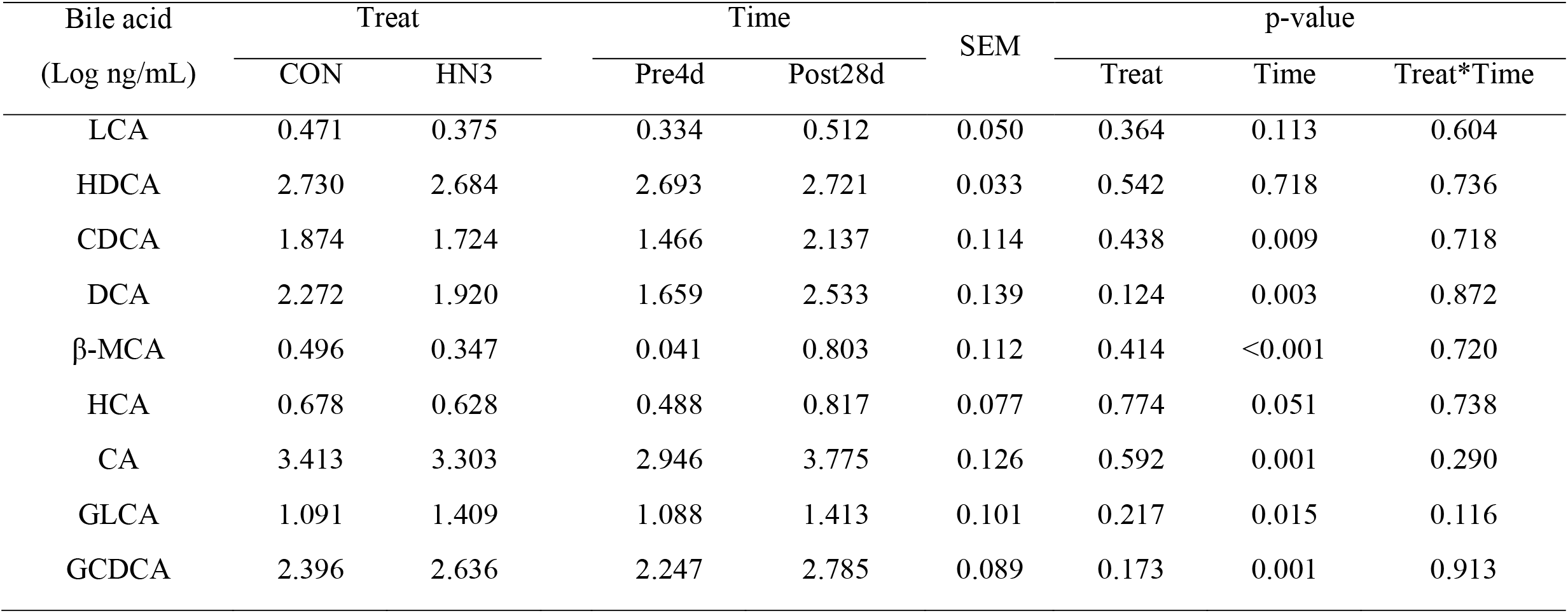
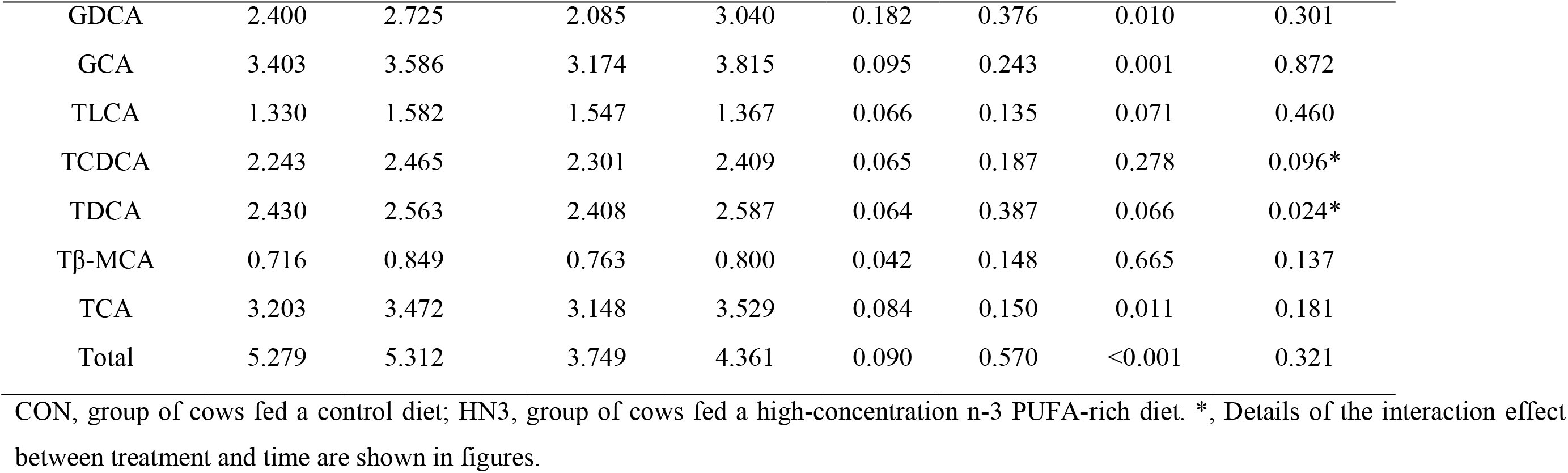
Bile acid concentration in the serum.

| Bile acid<br>(Log ng/mL) | Treat |  | Time |  | SEM | p-value |  |  |
| --- | --- | --- | --- | --- | --- | --- | --- | --- |
|  | CON | HN3 | Pre4d | Post28d |  | Treat | Time | Treat*Time |
| LCA | 0.471 | 0.375 | 0.334 | 0.512 | 0.050 | 0.364 | 0.113 | 0.604 |
| HDCA | 2.730 | 2.684 | 2.693 | 2.721 | 0.033 | 0.542 | 0.718 | 0.736 |
| CDCA | 1.874 | 1.724 | 1.466 | 2.137 | 0.114 | 0.438 | 0.009 | 0.718 |
| DCA | 2.272 | 1.920 | 1.659 | 2.533 | 0.139 | 0.124 | 0.003 | 0.872 |
| β-MCA | 0.496 | 0.347 | 0.041 | 0.803 | 0.112 | 0.414 | <0.001 | 0.720 |
| HCA | 0.678 | 0.628 | 0.488 | 0.817 | 0.077 | 0.774 | 0.051 | 0.738 |
| CA | 3.413 | 3.303 | 2.946 | 3.775 | 0.126 | 0.592 | 0.001 | 0.290 |
| GLCA | 1.091 | 1.409 | 1.088 | 1.413 | 0.101 | 0.217 | 0.015 | 0.116 |
| GCDCA | 2.396 | 2.636 | 2.247 | 2.785 | 0.089 | 0.173 | 0.001 | 0.913 |
| GDCA | 2.400 | 2.725 | 2.085 | 3.040 | 0.182 | 0.376 | 0.010 | 0.301 |
| GCA | 3.403 | 3.586 | 3.174 | 3.815 | 0.095 | 0.243 | 0.001 | 0.872 |
| TLCA | 1.330 | 1.582 | 1.547 | 1.367 | 0.066 | 0.135 | 0.071 | 0.460 |
| TCDCA | 2.243 | 2.465 | 2.301 | 2.409 | 0.065 | 0.187 | 0.278 | 0.096* |
| TDCA | 2.430 | 2.563 | 2.408 | 2.587 | 0.064 | 0.387 | 0.066 | 0.024* |
| Tβ-MCA | 0.716 | 0.849 | 0.763 | 0.800 | 0.042 | 0.148 | 0.665 | 0.137 |
| TCA | 3.203 | 3.472 | 3.148 | 3.529 | 0.084 | 0.150 | 0.011 | 0.181 |
| Total | 5.279 | 5.312 | 3.749 | 4.361 | 0.090 | 0.570 | <0.001 | 0.321 |

CON, group of cows fed a control diet; HN3, group of cows fed a high-concentration n-3 PUFA-rich diet. \*, Details of the interaction effect between treatment and time are shown in figures.

**Table 7.** Bile acid concentration in the liver.

| Bile acid<br>(Log ng/g) | Group |  | Time |  | SEM | p-value |  |  |
| --- | --- | --- | --- | --- | --- | --- | --- | --- |
|  | CON | HN3 | Pre4d | Post28d |  | Group | Time | Group*Time |
| LCA | 2.049 | 1.537 | 1.513 | 2.082 | 0.105 | 0.027 | 0.017 | 0.085* |
| HDCA | 3.917 | 4.410 | 4.569 | 3.758 | 0.154 | 0.105 | 0.032 | 0.096* |
| CDCA | 2.610 | 2.632 | 2.538 | 2.706 | 0.104 | 0.933 | 0.515 | 0.881 |
| DCA | 2.740 | 3.346 | 3.113 | 2.964 | 0.143 | 0.151 | 0.565 | 0.281 |
| ω-MCA | 1.338 | 1.182 | 1.037 | 1.484 | 0.081 | 0.332 | 0.046 | 0.609 |
| HCA | 3.347 | 3.342 | 3.345 | 3.345 | 0.019 | 0.904 | 0.987 | 0.690 |
| CA | 3.459 | 3.147 | 3.063 | 3.543 | 0.088 | 0.092 | 0.032 | 0.222 |
| GLCA | 0.946 | 0.904 | 0.837 | 1.020 | 0.064 | 0.701 | 0.197 | 0.386 |
| GCDCA | 2.907 | 2.821 | 2.758 | 2.961 | 0.057 | 0.459 | 0.120 | 0.610 |
| GDCA | 3.251 | 3.053 | 3.034 | 3.256 | 0.061 | 0.190 | 0.124 | 0.975 |
| GCA | 4.853 | 4.778 | 4.606 | 5.025 | 0.069 | 0.547 | 0.032 | 0.622 |
| TLCA | 2.577 | 2.555 | 2.626 | 2.505 | 0.031 | 0.649 | 0.073 | 0.859 |
| TCDCA | 3.459 | 3.479 | 3.534 | 3.398 | 0.052 | 0.943 | 0.240 | 0.283 |
| TDCA | 3.780 | 3.744 | 3.829 | 3.687 | 0.069 | 0.798 | 0.299 | 0.271 |
| TCA | 4.788 | 4.847 | 4.866 | 4.766 | 0.056 | 0.696 | 0.456 | 0.619 |
| Total | 5.279 | 5.312 | 5.267 | 5.324 | 0.041 | 0.725 | 0.559 | 0.513 |

CON, group of cows fed a control diet; HN3, group of cows fed a high-concentration n-3 PUFA-rich diet. The full forms of bile acid names are mentioned in Table S1.\*, Details of the interaction effect between treatment and time are shown in pictures.

#### Effect of time on the concentration of bile acids

The concentration of most BAs in serum increased from day 4 before calving to day 28 after calving (CDCA, *P* < 0.01; DCA, *P* < 0.01; ß-MCA, *P* < 0.001; HCA, *P* = 0.051; CA, *P* < 0.01; GLCA, *P* < 0.05; GCDCA, *P* < 0.01; GDCA, *P* < 0.05; GCA, *P* < 0.01; TCA, *P* < 0.05; total BAs, *P* < 0.01) (Table 6). However, the concentration of TLCA (*P* = 0.071) in serum decreased from day 4 before calving to day 28 after calving (Table 6). The LCA, ω-MCA, CA and GCA in the liver increased from day 4 before calving to day 28 after calving, but the HDCA in the liver decreased from day 4 before calving to day 28 after calving (Table 7).

### Liver transcriptomic analysis

#### Effect of diet on differential gene expression

DEGs were identified based on the threshold of *P*-values of <0.05 and |log2 fold change| values of >2. A total of 36 and 67 DEGs were identified between the CON and HN3 groups during the pre- and post-calving periods, respectively (Table 8).

**Table 8.** Effect of diet and time on differential gene expression.

| Comparison | | $0.01 \leq P < 0.05$ | $0.001 \leq P < 0.01$ | $P < 0.001$ |
| --- | --- | --- | --- | --- |
| Pre4d<br>HN3vsCON | Total | 25 | 8 | 3 |
|  | Up | 12 | 4 | 3 |
|  | down | 13 | 4 | 0 |
| Post28d<br>HN3vsCON | Total | 41 | 19 | 7 |
|  | Up | 11 | 5 | 3 |
|  | down | 30 | 14 | 4 |
| CON | Total | 50 | 37 | 30 |
| Post28d vs Pre4d | Up | 31 | 10 | 7 |
|  | down | 19 | 27 | 23 |
| HN3 | Total | 51 | 18 | 32 |
|  | Up | 30 | 8 | 17 |
| Post28d vs Pre4d | down | 21 | 10 | 15 |

CON, group of cows fed a control diet; HN3, group of cows fed a high-concentration n-3 PUFA-rich diet.

#### Effect of time on differential gene expression

A total of 117 DEGs (|log2 fold change| >2, *P* < 0.05) were identified between the pre-calving (4 days before calving) and post-calving (28 days after calving) periods in the CON group. In addition, 101 DEGs (log2 fold change| >2, *P* < 0.05) were identified between the pre-calving (4 days before calving) and post-calving (28 days after calving) periods in the HN3 group (Table 8).

#### Effects of diet on the results of KEGG pathway analysis

The results of KEGG pathway analysis of the top 20 DEGs between the CON and HN3 groups are shown in Figures S4–S5. These DEGs primarily participated in pathways related to taste transduction during the pre- and post-calving periods. Therefore, taste transduction-related genes were the main focus of further analysis. Genes involved in taste transduction are listed in Table 9. Additionally, these DEGs participated in pathways associated with lipid metabolism, such as regulation of lipolysis in adipocytes and biosynthesis of unsaturated fatty acids, during the pre-calving period and in pathways associated with herpes simplex virus infection and circadian rhythm during the post-calving period.

**Table 9.**
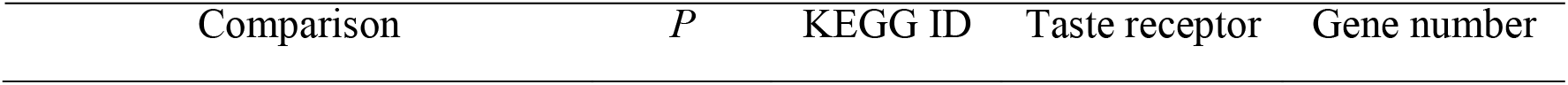
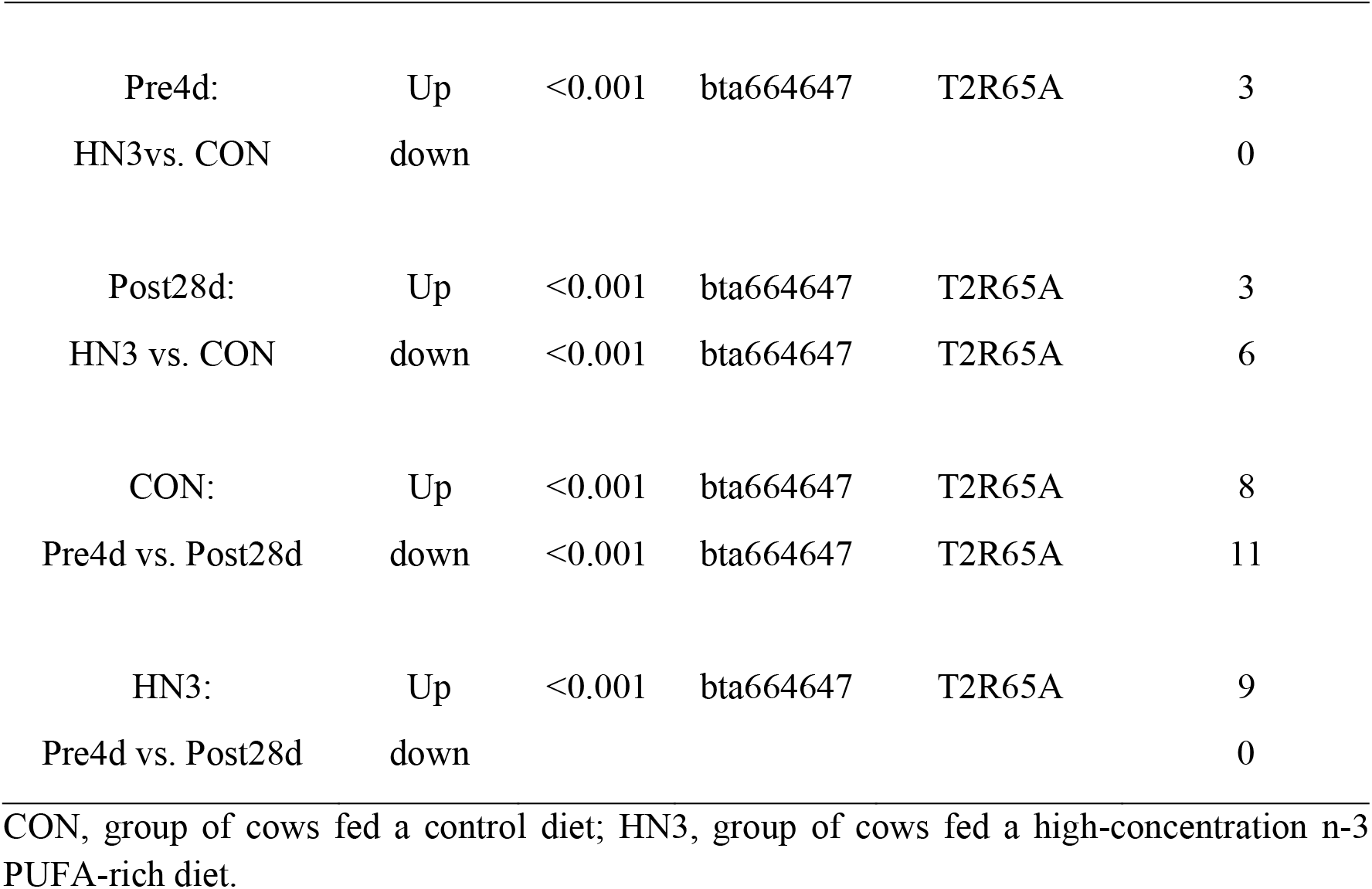
KEGG analysis of differentially expressed genes in the liver.

| Comparison | *P* | KEGG ID | Taste receptor | Gene number |
| --- | --- | --- | --- | --- |

|  |  |  |  |  |  |
| --- | --- | --- | --- | --- | --- |
| Pre4d: | Up | <0.001 | bta664647 | T2R65A | 3 |
| HN3vs. CON | down |  |  |  | 0 |
| Post28d: | Up | <0.001 | bta664647 | T2R65A | 3 |
| HN3 vs. CON | down | <0.001 | bta664647 | T2R65A | 6 |
| CON: | Up | <0.001 | bta664647 | T2R65A | 8 |
| Pre4d vs. Post28d | down | <0.001 | bta664647 | T2R65A | 11 |
| HN3: | Up | <0.001 | bta664647 | T2R65A | 9 |
| Pre4d vs. Post28d | down |  |  |  | 0 |

CON, group of cows fed a control diet; HN3, group of cows fed a high-concentration n-3 PUFA-rich diet.

#### KEGG pathway analysis before and after calving

Similarly, KEGG analysis of DEGs identified between the pre- and post-calving periods revealed that the DEGs were enriched in pathways associated with taste transduction in the HN3 and CON groups. In addition, these DEGs were highly enriched in pathways associated with African trypanosomiasis, dilated cardiomyopathy and IgA production (Figures S6–S7).

### Correlation analysis

The correlation between DMI and different parameters, such as the concentration of acetate, propionate, butyrate and total VFAs; the acetate-to-propionate ratio (A:P); the relative bacterial abundance and the expression of taste transduction-related genes, is shown in Figure 2. The DMI of cows was positively correlated with the concentration of orexins (r = 0.742, *P* < 0.001) and NPY (r = 0.687, *P* < 0.001) in serum and BAs in the liver (HDCA: r = 0.501, *P* < 0.05; MCA: r = 0.486, *P* < 0.05; GCA: r = 0.502, *P* < 0.05). In addition, DMI was positively correlated with the relative abundance of *Spirochaetota* (r = 0.871, *P* < 0.001) and *Bacteroidota* (r = 0.896, *P* < 0.001) in the rumen and the expression of genes involved in taste transduction (ACSL1: r = 0.673, *P* < 0.001; PLIN4: r = 0.632, *P* < 0.01; CPT1A: r = 0.694, *P* < 0.001; ANGPTL4: r = 0.479, *P* < 0.05; APOA1: r = 0.813, *P* < 0.001; RXRG: r =0.447, *P* < 0.05; PCK1: r = 0.559, *P* < 0.01). However, DMI was negatively correlated with the concentration of VFAs (acetate: r = -0.652, *P* < 0.001; A:P ratio: r = -0.640, *P* < 0.01); the concentration of LCA in the liver (r =-0.827, *P* < 0.001) and the abundance of *Desulfobacteria* (r = -0.479, *P* < 0.05), *Elusimicrobiota* (r = -0.614, *P* < 0.05), *WPS-2* (r = - 0.426, *P* < 0.05) and *Actinobacteriota* (r = -0.573, *P* < 0.01) in the rumen.

### QPCR validation on gene expression

The relative expression of ACSL1 gene (*P* < 0.01) and PLIN4 gene (*P* < 0.05) were upregulated in HN3 than that in CON during pre-calving and post-calving period (Figure 3). The relative expression of CPTIA was higher in HN3 than CON during pre-calving (*P* < 0.05), but not affected by diet during post-calving (Figure 3). There was no difference for ACSL1 and PLIN4 gene expression in liver from 4 d pre-calving to 28 d post-calving. However, the CPTIA gene expression was higher in 4 d pre-calving than in 28 d post-calving (*P* < 0.01).

## Discussion

### Effect of n-3 PUFA on DMI

Milk production during the post-calving period was increased by 6.25 kg in the HN3 group. Simultaneously, compared with the control diet, the diet supplemented with high n-3 PUFA significantly increased the DMI of cows during the transition period. These results are consistent with those of studies by Petit et al. (2007) and Zachut et al. (2010), who reported that cows (transition period) fed a diet rich in saturated FAs had a lower milk yield and DMI than those fed a diet rich in unsaturated FAs. However, Gonthier et al. (2005) reported that feeding extruded flaxseed enriched in n-3 PUFA at 12.7% of DM did not alter the DMI of lactating cows (225 days after calving), whereas Chilliard et al. (2009) reported that feeding extruded flaxseed at 21.2% of DM decreased the DMI of cows (213 days after calving). These inconsistent results may be related to differences in the concentration of supplements and the status of cows or the interaction of dietary supplements with other dietary components Chilliard et al. (2009). Increasing the dietary intake of n-3 PUFA through the addition of extruded flaxseed may increase DMI during the transition period but not during the lactation period. In this study, n-3 PUFA supplementation increased DMI, resulting in a dramatic increase in milk production. This increase in DMI may be associated with various factors including fermentation parameters, bacterial abundance and gene expression.

### Effect of n-3 PUFA on rumen fermentation

The pH and concentration of VFAs are essential characteristics for assessing fermentation in the rumen (Liu et al., 2012). In this study, feeding n-3 PUFA to cows did not affect ruminal pH and the concentration of VFAs during the transition period. This result was consistent with that of previous studies on lactating dairy cows fed diets containing 10% (wt/wt) of crushed flaxseeds (Beauchemin et al., 2009) and on sheep fed diets containing linseed oil (Ueda et al., 2003, Kim et al., 2007). However, a study reported that a diet supplemented with 90 g of linseed oil/d decreased the concentration of total VFAs (Czerkawski et al., 1975). These inconsistent results may be attributed to differences in the amount and form of n-3 PUFA supplementation. However, in this study, the concentration of acetate and the acetate-to-propionate ratio were found to decrease from the pre-calving to the post-calving period. This change can be explained by the low ratio of forage-to-concentrate in the diet during the post-calving period; the increased dietary concentrate feed level decreased the molar ratio of acetate-to-propionate (Liu et al., 2019). In addition, DMI was negatively correlated with the concentration of acetate and the acetate-to-propionate ratio, which is consistent with the results of a study by Seymour et al. (2005).

### DMI regulation with plasma biomarkers

Regulation of dietary intake is a complex biological phenomenon. Eating behaviour and food intake are the results of neural integration of numerous signals related to the environment, feed and physiological state of the animal. Some intake-stimulating neuropeptides such as NPY and orexins play an important role in mediating the effects of peripheral signals to the brain and are potentially involved in intake regulation (Chen et al., 2012). In this study, a diet supplemented with extruded flaxseed (n-3 PUFA) increased the concentration of NPY and orexins in serum. This increase is associated with the increase in DMI owing to n-3 PUFA supplementation during the transition period. These results are inconsistent with those of previous studies, which have demonstrated that n-3 PUFA supplementation in rats effectively promoted weight loss by reducing DMI and the expression of NPY (Jia et al., 2009, Ma et al., 2016). A possible reason underlying these inconsistent results is as follows: Microorganisms in the rumen play an important role in n-3 PUFA catabolism and indirectly affect the expression of NPY in cows during the transition period. A study demonstrated that treatment with a mixture of *Bifidobacterium lactis*, *Lactobacillus rhamnosus* GG and inulin altered the gut microflora and elevated the portal plasma levels of NPY (Lesniewska et al., 2006).

### DMI regulation with ruminal bacteria

Ruminal bacteria play an important role in digesting complex and simple carbohydrates, and their abundance is associated with the DMI of cows. The fermentation of feed organic matter by ruminal microbes and microbial biomass synthesis can meet 70–85% of the energy requirements and 70–100% of the protein requirements of ruminants (Thirumalesh and Krishnamoorthy, 2013). In this study, n-3 PUFA supplementation in cows increased the relative abundance of *Spirochaetota* during the pre- and post-calving periods, *Bacteroidota* during the pre-calving period and *Elusimicrobia*. The abundance of *Spirochaetota* is associated with extensive carbohydrate digestion and metabolism in the camel rumen (Gharechahi et al., 2022). In addition, *Spirochaetota* may contribute to increase cellulose- and hemicellulose-degrading enzymes activities (Gharechahi and Salekdeh, 2018). *Bacteroidota* is a dominant phylum in the rumen, accounting for approximately half of the total bacterial abundance. *Bacteroidota* can potentially degrade a wide range of carbohydrate polymers and produce various VFAs (Stewart et al., 2018, Stewart et al., 2019). According to the function of these bacteria, n-3 PUFA supplementation should increase the concentration of VFAs in the rumen. However, this phenomenon was not observed in this study. This discrepancy may be attributed to the complex processes of rumen fermentation and VFA production, both of which involve the interaction between diverse microorganisms, and is related to the absorption ability of the rumen wall (da Silva ALONSO et al., 2006). In this study, we did not test the rumen wall absorption ability, which should be investigated in future studies.

The abundance of *Elusimicrobia* is negatively correlated with urinary N excretion (Alves et al., 2021). This phylum has been detected in the intestine of termites (Geissinger et al., 2009) and ruminants (Deusch et al., 2017); however, it is not involved in the digestion of plant fibres in the gut and functions through an unusual peptide degradation pathway involving transamination reactions (Herlemann et al., 2009). In this study, the concentration of NH_3_-N was positively correlated (r = 0.745, *P* < 0.001) with the relative abundance of *Elusimicrobia*, suggesting the involvement of *Elusimicrobia* in protein catabolism in the rumen.

### DMI regulation with liver BAs

*De novo* biosynthesis of BAs (primary BAs), such as CA and CDCA, occurs in the liver. An important function of the liver is the secretion of BAs into bile and their subsequent release into the duodenum (Grüner and Mattner, 2021). Approximately 15% of BAs escape the absorption of the terminal ileum and migrate to the colon, where gut microbes promote the biotransformation of primary BAs to secondary BAs, such as DCA, LCA and UDCA (Di Ciaula et al., 2018). Some BAs are re-absorbed in the terminal ileum and colon and return to the liver via the portal vein (Di Ciaula et al., 2018). BAs are involved in several important functions in the liver and intestine. They facilitate the elimination of cholesterol from the liver into bile and promote the absorption of lipids and lipid-soluble vitamins from the intestine (Li and Lu, 2018). However, accumulation of some secondary BAs (DCA and LCA) in the liver may cause damage to the liver and is negatively correlated with DMI. A study showed that exacerbated BA metabolism induced by low feed intake significantly increased the levels of toxic BAs, DCA and LCA, in piglets during the transition period (Lin et al., 2018). This finding may at least partially explain how n-3 PUFA supplementation-induced decrease in LCA concentration in the liver was associated with an increase in DMI during the post-calving period. Consistently, correlation analysis revealed that LCA concentration in the liver was strongly negatively correlated with DMI.

### DMI regulation with liver DEGs

DEGs identified from the liver tissues of cows between the CON and HN3 groups during the pre- and post-calving periods were significantly enriched in the taste transduction-related pathway (bta04742), which involves the taste receptor T2R65A. The proteins encoded by these genes are co-expressed in distinct subpopulations of taste bud cells of the human gustatory system (Brunes et al., 2021). Although taste perception affects feed intake and production traits (Brunes et al., 2021), genes involved in the taste transduction-related pathway are rarely reported as candidate genes for feed intake in cattle. In this study, the expression of some genes involved in the taste transduction-related pathway (e.g. ACSL1, PLIN4, CPTIB, GK and CPTIA) was positively correlated with DMI. These genes play a role in fatty acid metabolism. For instance, ACSL participates in fatty acid metabolism by converting fatty acids to fatty acyl-CoAs to regulate various physiologic processes. Perilipin family (PLIN) members can facilitate the movement of lipid droplets (Han et al., 2018), which are functional subcellular organelles involved in lipid metabolism (Chen et al., 2013). The CPT1B gene regulates lipid metabolism in Chinese Simmental cattle (Schlaepfer and Joshi, 2020) and may prevent fatty liver disease (Dang et al., 2020). Both CPT1A and CPT1B are involved in the oxidation of long-chain fatty acids and nucleotide biosynthesis (Schlaepfer and Joshi, 2020). GK participates in impaired insulin suppressibility of hepatic glucose production in rats fed a high-fat diet (Oakes et al., 1997). The expression of the abovementioned taste transduction-related genes, which are also involved in fatty acid metabolism, may be regulated in cows transitioning from the pre- to post-calving period. Our study suggests that dietary supplementation with n-3 PUFA can upregulate the expression of ACSL1, CPTIA and PLIN4, resulting in an increase in DMI.

## Conclusion

Dietary supplementation with n-3 PUFA significantly increased milk production during the post-calving period by increasing DMI during the pre- and post-calving periods. DMI was positively correlated with the concentration of NPY and orexins in serum. n-3 PUFA may affect the expression of genes involved in taste transduction and subsequently decrease the concentration of toxic BAs (such as LCA) in the liver. In addition, DMI was positively correlated with the abundance of *Spirochaetota*, *Bacteroidota* and *Elusimicrobia* in the rumen of cows. However, mechanisms underlying the effects of these factors on each other could not be examined owing to the limitation of phylogenetic resolution. We speculate that dietary supplementation with n-3 PUFA at an appropriate concentration can benefit the milk production and DMI of cows during the transition period, and taste transduction-related genes play an important role in this beneficial effect.

## Author Contributions

Conceptualization, X.S., S.L. and W.W.; methodology, W.W., and Z.C; software, X.S.; validation, S.L.; formal analysis, X.S.; investigation, X.S., Z.Y., Y.Z., Q.W., Z.W., C.G and T.X.; resources, C.G.; data curation, X.S.; writing—original draft preparation, X.S.; writing—review and editing, W.W.; visualization, X.S.; supervision, S.L.; project administration, W.W.; and funding acquisition, S.L. All authors have read and agreed to the published version of the manuscript.

## Funding

This research was funded by the National Natural Science Foundation of China (32130100), and The 2115 Talent Development Program of China Agricultural University.

### Institutional Review Board Statement

All experimental procedures were approved by the China Agricultural University Laboratory Animal Welfare and Animal Experimental Ethical Inspection Committee.

### Data Availability Statement

The rumen microbial16S RNA and liver transcriptome sequencing data were uploaded to NCBI (16s RNA project: PRJNA913816; transcriptome project: PRJNA915484). Other data that support the findings of this study are available from the corresponding author upon reasonable request.

## Acknowledgments

The authors thank Zhongdi Animal Husbandry Technology Company for providing the experimental animals.

